# Reduced processing and toxin binding associated with resistance to Vip3Aa in a resistant strain of fall armyworm (*Spodoptera frugiperda*) from Louisiana

**DOI:** 10.1101/2025.02.27.640590

**Authors:** Rajeev Roy, Heba Abdelgaffar, Dawson Kerns, Matthew Huff, Margaret Staton, Fei Yang, Fangneng Huang, Juan Luis Jurat-Fuentes

## Abstract

**Background:** Transgenic crops expressing Cry and Vip3Aa insecticidal proteins from the bacterium *Bacillus thuringiensis* are a primary tool for controlling fall armyworm (*Spodoptera frugiperda*) populations. The evolution of resistance to Cry proteins in the native range of the fall armyworm has increased reliance and intensified the selection of resistance to Vip3Aa. In this study, we identified mechanisms of resistance to Vip3Aa in the LA-RR strain of *S. frugiperda* originating from Louisiana (USA).

**Results:** Midgut epithelial damage in susceptible larvae was evidenced by a significant drop in midgut pH after feeding on either Vip3Aa protoxin or activated toxin. In contrast, this midgut pH drop was only detected for activated Vip3Aa toxin in LA-RR larvae. Midgut fluids from LA-RR larvae displayed delayed processing of Vip3Aa protoxin when compared to fluids from susceptible larvae, and this slower processing was associated with reduced activity and expression of trypsin and chymotrypsin enzyme genes in the LA-RR strain. In bioassays, LA-RR larvae were significantly more susceptible to Vip3Aa protoxin pre-processed by midgut fluids from susceptible than from LA-RR larvae. In addition, midgut brush border membrane vesicles from LA-RR larvae exhibited lower specific Vip3Aa toxin binding than vesicles from the susceptible strain.

**Conclusion:** The results of this study support that both slower proteolytic processing and reduced specific binding are associated with resistance to Vip3Aa in an *S. frugiperda* strain from the Western hemisphere, the native range of this pest. This information increases our understanding of resistance to Vip3Aa and advances monitoring and fall armyworm management.

## 1 INTRODUCTION

Larvae of the fall armyworm (FAW, *Spodoptera frugiperda*) are one of the most damaging pests of corn and other crops, and the distribution and economic impact of this pest are expanding dramatically from its native range in the Western Hemisphere to threaten global food security.^1^ Transgenic plants expressing Cry and/or Vip3Aa insecticidal proteins from the bacterium *Bacillus thuringiensis* (Bt crops) provide control of FAW populations in agricultural settings.^2,3^ In the Americas, transgenic corn and cotton expressing the Cry1F protein were commercialized primarily to control FAW.^4,5^ However, practical FAW resistance to Cry1F evolved in Puerto Rico,^6^ Florida,^7,13^ Brazil,^8^ and Argentina.^9^ Shared binding sites for Cry1A and Cry1F proteins in the midgut of FAW larvae^10,11^ explained cases of cross-resistance to these proteins observed in FAW populations resistant to at least one Cry1A or Cry1F protein.^12-17^

Reduced efficacy of Cry proteins against some agriculturally important pests has augmented reliance on Vip3Aa traits,^18^ further increasing the pressure for the evolution of resistance. Although field failures from resistance to Vip3Aa have not yet been documented for FAW, multiple resistance alleles have been isolated from field-collected samples in the U.S., China, and Brazil.^19-23^ Moreover, available estimates support a relatively high frequency of Vip3Aa-resistance alleles in field FAW populations,^14^ violating assumptions in current resistance management approaches for Bt crops.^24^

Resistance can emerge from the alteration of any step in the Vip3Aa mode of action, which is not completely understood.^25^ After ingestion, non-toxic Vip3Aa protoxin is processed by midgut serine proteases to an active Vip3Aa toxin form through a drastic structural change.^26, 27^ The Vip3Aa toxin interacts in an unknown manner with chitin on the midgut peritrophic matrix, and this interaction seems critical for toxicity.^28^ Subsequently, the Vip3Aa toxin binds to specific receptors on the insect midgut epithelium, which are not recognized by Cry proteins.^29,30^ Binding is followed by Vip3Aa pore formation and cell death by osmotic shock.^31,32^ There is also evidence for Vip3Aa internalization through endocytosis and subsequent apoptosis, but only from studies with insect cell cultures.^33,34^

While relatively few studies are available, altered processing of protoxin to toxin is the most common mechanism of resistance to Vip3Aa in Lepidoptera,^35,36^ with evidence of reduced Vip3Aa toxin binding associated with resistance reported only in a strain of *Helicoverpa zea*.^37^ In FAW so far, resistance to Vip3Aa has been linked to downregulation of the *SfMyb* transcription factor^22^ and loss of chitin synthase-2 function^23^ in two distinct laboratory-selected strains from China, although the specific resistance mechanisms were not identified. Results from functional tests identify the expression of chitin-synthase 2 and the *SfVipR1* gene as critical for susceptibility to Vip3Aa.^23,38^ However, chitin-synthase 2 did not function as a Vip3Aa receptor when expressed in cultured insect cells.^23^ So far, there is no information on mechanisms of resistance to Vip3Aa in field populations from the Western Hemisphere, the native range of FAW.

The present study aimed to determine the mechanism of monogenic and autosomal resistance to Vip3Aa in a strain of FAW (LA-RR) generated from field collections in Louisiana (USA).^19^ None of the *SfMyb, SfVipR1*, and chitin synthase-2 alterations described in Vip3Aa-resistant FAW strains from China were associated with resistance in LA-RR, supporting that it may harbor a novel mechanism of resistance.

## 2 MATERIALS AND METHODS

### 2.1 Insect colonies

The LA-SS strain of FAW was established in 2016 from larvae collected from non-Bt corn fields in Franklin Parish (Louisiana, USA).^39^ This strain has been continuously reared in the laboratory without pesticide exposure and remains susceptible to Cry1F, Cry1A.105, Cry2Ab2, Cry2Ae, and Vip3Aa.^40^ During the performance of this study, the LA-SS unexpectedly collapsed. Consequently, further work was performed using the Benzon strain of FAW as the susceptible reference strain, as it displays Cry1F and Vip3Aa susceptibility similar to LA-SS.^12,41^ Eggs of this strain were purchased from Benzon Inc. (Carlisle, PA, USA).

The LA-RR strain of FAW was developed in 2016 from an F_2_ screen with leaf tissue of Viptera 3111 corn producing the Vip3Aa20 protein of field-collected larvae from Franklin and Rapides parishes in Louisiana.^19^ Surviving (resistant) larvae were reared under selection with Viptera 3111 corn leaf tissue for more than six generations, followed by selection with a discriminatory concentration (3.16 μg/cm^2^) of purified Vip3Aa39 protoxin.^19^ Larvae of the LA-RR strain remain susceptible to Cry1F, Cry1A.105, Cry2Ab2, and Cry2Ae but are highly resistant (>632-fold) to Vip3Aa protoxin.^19,42^

### 2.2 Transcriptome profiling

Total RNA was extracted individually from dissected midguts of 4^th^ instar FAW larvae from the LA-SS and LA-RR strains (six midgut samples per strain) using the Direct-Zol RNA miniprep kit (Zymo Research, Irvine, CA, USA), following the manufacturer’s instructions. The extracted RNA was quantified and purity examined using UV-VIS absorbance at 260/230 nm and 260/320 nm in a Nanodrop-One spectrophotometer (Thermo-Fisher Scientific, Waltham, MA, USA). The total RNA samples were then sent for mRNA enrichment, library construction, and Illumina short-read sequencing on a 150 bp paired-end read configuration at Novogene (Sacramento, CA, USA). The raw transcript sequences were deposited in the National Center for Biotechnology Information Sequence Read Archive (NCBI SRA) database as accession no. PRJNA1196488.

Raw reads were first quality checked with fastQC ^43^ and then the low-quality bases and adaptors were removed by Skewer.^44^ The cleaned reads were then mapped to the reference *S. frugiperda* genome assembly (AGI-APGP_CSIRO_Sfru_2.0, accession no: GCF_023101765.2)^45^ using STAR.^46^ The resultant bam files were analyzed by Htseq-count to generate raw read counts for all genes,^47^ which were analyzed in R-studio using the DESeq2 package to estimate the total number of differentially expressed genes between the strains.^48^ Genes that displayed consistent regulation trends across biological replicates (eg: downregulated in all resistant and upregulated in all susceptible replicates), had a log2Fold change >1, and P-adjusted < 0.05 were considered further. The list of upregulated and downregulated genes that met this threshold was analyzed for functional groups by GhostKoala.^49^

### 2.3 Quantitative PCR

The expression of selected genes was verified with real-time quantitative PCR (RT qPCR) as previously described.^50^ Total RNA was extracted from individual midguts as described for RNAseq, and then 1 µg of total RNA was used to synthesize first-strand cDNA using the SuperScript III reverse transcriptase kit (Invitrogen, Waltham, MA, USA), following the manufacturer’s instructions. Specific primers for each transcript were designed using the Primer-Blast tool.^51^ Quantitative PCR was conducted in 10 µL reactions containing 5 µL of SYBR Green 2X Master Mix (Applied Biosystems, Foster City, CA, USA), 3 µL of cDNA (20 ng), 1 µL of each primer (10 µM stocks), and 2 µL of water in a QuantStudio 6 Flex Real-Time PCR System (Applied Biosystems). The expression analyses were performed using the 2^-ΔΔCT^ method^52^ with the ribosomal L-17 gene (XM_035591646.2) as the internal reference.^53^ The relative expression in the graphical representation of the data was normalized to consider the average expression in LA-SS as 1. The relative efficiency of primers was estimated from standard curves with primer dilutions and the LA-SS cDNA as a template. Primer sequences and efficiencies for the three target genes are listed in Supporting Information Table S1.

### 2.4 Quantification of midgut chitin

Chitin was quantified in dissected midguts of 4^th^ instar FAW larvae as described in Henriques et al.^54^ Individual dissected midguts from actively feeding larvae with an intact food bolus were homogenized in 300 μL of distilled water with a TissueLyser II sample disruptor (QIAGEN, Germantown, MD, USA) at 30 Hz for 5 min. Samples were clarified with a short spin cycle of 3 seconds, and 100 μL of the supernatant was transferred to a new 1.5 mL microfuge tube containing 100 μL of calcofluor stain solution (Sigma Aldrich, St Louis, MO, USA) and 300 μL of distilled water. After incubation for 15 minutes in the dark, the samples were centrifuged at 21,000 x *g* for 5 minutes, and the supernatants were discarded. The pellets were submitted to washing with 200 μL of water and centrifugation steps twice, and the final pellets were resuspended in 100 μL of distilled water, and then fluorescence at 355 nm was measured in a Biotek Synergy H1 microplate reader (Agilent, Santa Clara, Ca, USA) for 1 μL of each sample in 100 μL of distilled water in a 96-well microplate (1/2 Area OptiPlate-96, Revvity, Waltham, MA, USA). Mean chitin amounts and corresponding standard errors were calculated from determinations of 12 larval midguts from each strain. Significant differences between fluorescent values were tested by pooled t-test (P < 0.05) using GraphPad PRISM 10.2.3 for Windows (GraphPad Software, Boston, MA, USA).

### 2.5 Vip3Aa protein

A recombinant *Escherichia coli* strain harboring the *vip3Aa39* gene was used to produce the Vip3Aa39 protein following procedures described elsewhere,^37^ except that when purifying Vip3Aa39 for radiolabeling 0.1% β-mercaptoethanol was added to the final lysis buffer. This addition was found to reduce over-iodination of the toxin during radiolabeling. The Vip3Aa39 protein has >95% sequence identity to the Vip3Aa19 and Vip3Aa20 traits produced in Bt corn and cotton, respectively, and shows indistinguishable insecticidal activity to the Vip3Aa19 protein against *H. zea*.^37^ Based on this high identity and similar activity, we refer to the Vip3Aa39 protein used in this study as Vip3Aa.

The Vip3Aa protoxin and the toxin form obtained after processing protoxin with trypsin for 2 hours at 37°C were purified using anion exchange chromatography in a HiTrap Q HP (Cytiva, Marlborough, MA, USA) column equilibrated in buffer A (50 mM Na_2_CO_3_, 50 mM NaHCO_3,_ pH 9.8) connected to an AKTA Pure FPLC (GE Healthcare, Uppsala, Sweden). The fractions eluted with a linear gradient of buffer B (buffer A with 1M NaCl) were resolved by SDS-10%PAGE, and proteins in gels were stained with ProtoBlue Safe (National Diagnostics, Atlanta, GA, USA) to identify fractions containing Vip3Aa based on the expected molecular size of 88-kDa (protoxin) and 66-kDa (toxin). Fractions containing purified Vip3Aa protoxin or toxin were pooled, and their concentration was estimated using the Qubit Protein Assay Kit (Invitrogen) before storing at -80°C until used.

### 2.6 *In situ* measurements of gut pH

The pH in sections of the digestive tube was measured as described elsewhere.^37,55^ Three cohorts of six actively feeding 4^th^ instar larvae per strain were starved overnight and then fed with a drop (10 µl) containing traces of bromophenol blue and 10% sucrose alone (control) or with of Vip3Aa39 protoxin (5 μg) or toxin (10 μg). After 2 hours of feeding, the larvae were immobilized and the dorsal cuticle cut open to expose the digestive tube. A micro pH electrode (Thermo Scientific, model 9863BN) was inserted to measure the pH in five different sections (anterior to posterior): foregut, anterior midgut, central midgut, posterior midgut, and hindgut. Only digestive tubes containing bromophenol blue traces as evidence of feeding were considered in the analyses. The significance of pH value differences for each treatment within a strain was tested with a one-way ANOVA (α = 0.05) using GraphPad Prism.

### 2.7 Gut fluid collections

Gut fluids were collected from actively feeding 4^th^ instar as described elsewhere^37,55^ from LA-SS, LA-RR, and Benzon larvae. Five dissected midguts were combined in a single microcentrifuge tube (considered one biological replicate) with deionized water (500 µL), and the contents were mixed well by vortexing before centrifugation at 15,000 rpm for 20 min at 4°C. The supernatant was carefully collected as the gut fluid sample in a new microcentrifuge tube, and the total protein concentration was measured by Qubit fluorometry (Invitrogen). The samples were diluted to 1 mg/mL with deionized water, aliquoted, and frozen at -80°C for two weeks.

### 2.8 Vip3Aa protoxin processing

The processing of purified Vip3Aa protoxin by midgut fluids from FAW larvae was performed as described previously,^37^ with minor adjustments. Based on the rate of processing observed in preliminary tests (data not shown), we selected 0.4 μg as the amount of gut fluid proteins to use in detecting and comparing the rate of Vip3Aa protoxin processing between FAW strains. Gut fluid proteins were mixed with 80 μg of Vip3Aa protoxin in a total volume of 140 µL of carbonate-bicarbonate buffer (50 mM Na_2_CO_3_, 50 mM NaHCO_3_, pH 9.8) and incubated at 37°C with constant stirring at 250 rpm. Aliquots (10 µL) were collected after 15 min, 30 min, 75 min, 90 min, and 120 min, and proteolytic processing was immediately stopped by adding 10 µL of 2X sample buffer^56^ and heat denaturing for 5 min at 95°C. All samples were then resolved on SDS-10%PAGE, and protein bands were visualized by staining with ProtoBlue Safe. An unprocessed Vip3Aa protoxin sample was included in the gel for reference. Processing of Cry1F protoxin (Supporting Information, Fig. S1) was performed as described for Vip3Aa protoxin.

### 2.9 Quantification of midgut soluble serine protease activity

Trypsin and chymotrypsin activities in midgut fluids were quantified using specific substrates as described elsewhere,^57^ with minor modifications. Trypsin activity was measured in triplicate in reactions containing 10 μL of gut fluids at a 1 μg/μL protein concentration and 10 µL (1 µg/µL stock) of trypsin substrate (L-BAPNA, Sigma Aldrich) in a final volume of 100 µL of carbonate-bicarbonate buffer. Processing of the substrate was monitored in 1 min intervals for 10 min by the release of ρ-nitroanilide and the corresponding change in absorbance at 405 nm using a Biotek Synergy H1 (Agilent) plate reader with the associated Gen5 software for data analysis. Chymotrypsin activity was quantified with the same reaction components as trypsin but using a fluorogenic substrate (Suc-Ala-Ala-Pro-Phe-AMC, Sigma Aldrich). Processing of the substrate was monitored in 30-sec intervals for 10 minutes by detecting fluorescence (Ex/Em= 380/460 nm) in the Biotek Synergy H1 plate reader. The significance of differences in the release of the products was tested using an unpaired t-test (α = 0.05) in GraphPad Prism.

### 2.10 Bioassays

The susceptibility of FAW larvae from the Benzon strain to Vip3Aa protoxin and toxin was estimated from surface overlay assays using a meridic diet (beet armyworm diet, Frontier Agricultural Sciences, Newark, DE, USA) in 128-well plastic bioassay trays (C-D International Inc, Pittman, NJ, USA). Solutions of Vip3Aa protoxin or toxin were overlaid on the surface diet of individual wells to obtain a range of toxin densities (0.01, 0.05, 0.1, 0.25, and 0.50 μg/cm^2^), and allowed to air dry. A single FAW neonate was placed per well, a total of 16 larvae were tested per treatment, and the bioassays were performed a total of four times. The bioassay trays were incubated at 18h L/ 6h D photoperiod and 26°C for seven days, and then mortality (lack of larval movement when prodded) was assessed.

A similar protocol was used for bioassays on LA-RR neonates with a discriminatory surface density of Vip3Aa protoxin (5 μg/cm^2^) based on previous estimates with LA-RR neonates.^59^ The purified Vip3Aa protoxin (140 μg) in these tests was pre-processed with 0.5 µg of midgut fluids from Benzon or LA-RR larvae in a total volume of 0.5 ml for two hours, as described for the *in vitro* protoxin processing assay, and mortality and body weight of survivors was assessed after nine days. Sixteen larvae were tested per sample, and the bioassay was performed a total of four times with eggs from two independent cohorts.

### 2.11 Brush border membrane vesicle (BBMV) preparation

Midguts were dissected from actively feeding 4^th^ instar *S. frugiperda* larvae and used to prepare BBMV using a differential magnesium precipitation method,^59^ with minor modifications.^37,60^ The protein concentration in the isolated BBMV was quantified by Qubit fluorometry and then they were stored at -80°C until used.

The quality of the BBMV preparations was estimated based on the enrichment in aminopeptidase (APN) activity as a brush border marker enzyme, as described elsewhere.^61^ Typical APN activity enrichment in BBMV samples compared to initial midgut homogenates ranged between 3 to 6-fold for independent BBMV preparations. There were no significant differences in total APN activity or enrichment between BBMV from susceptible and Vip3Aa-resistant strains (One Way ANOVA, Tukey Post Hoc test, P = 0.998).

### 2.12 Labeling of Vip3Aa

The purified Vip3Aa toxin was labeled with a 1:30 molar ratio of EZ-Link Sulfo-NHS-LC-Biotin (ThermoFisher) as described elsewhere.^37^ Free biotin was removed from the biotinylated samples by dialysis, and the labeled toxin was quantified using Qubit fluorometry. Successful biotinylation was confirmed by western blotting (data not shown).

Labeling of Vip3Aa toxin with Alexa Fluor-488 followed the manufacturer’s instructions (ThermoFisher). The labeled toxin was quantified using fluorometry (Qubit) and successful labeling (Supporting Information, Fig. S2A) and binding to midgut BBMVs (Supporting Information, Fig. S2B) were confirmed by resolving a labeled sample or BBMV with bound toxin by SDS-10%PAGE and then imaging at 488/525 nm (ex/em) on a ChemiDoc MP imaging system (Bio-Rad, Hercules, CA, USA) with the Pro Q emerald 488 filter.

Labeling of purified Vip3Aa toxin (25 μg) with NaI^125^ radioisotope (0.5 mCi, Revvity) was performed with chloramine T, as described previously.^37^ Specific activity for radiolabeled Vip3Aa ranged between 0.03 and 0.09 mCi/pmol.

### 2.13 BBMV binding assays

Western blotting testing Vip3Aa binding to BBMV was done as described elsewhere^37^ with biotinylated Vip3Aa (3 μg) and BBMV (30 μg) with or without a 100-fold molar excess of unlabeled Vip3Aa in a final reaction volume of 100 µL of binding buffer (137 mM NaCl, 10 mM Na_2_HPO_4_, 2.7 mM KCl, 1.8 mM KH_2_PO_4_, 0.1%BSA, 0.1% Tween-20, pH 7.4). After an hour at room temperature, reactions were stopped by centrifugation at 15,000 rpm for 15 minutes at room temperature. The supernatant was removed, and the pellet was resuspended in 500 µL of ice-cold binding buffer and centrifuged again. The final pellet was resuspended in 20 µL of 2X sample buffer,^56^ heat denatured at 95°C for 5 minutes, and then loaded on an SDS-10%PAGE gel. After electrophoresis, proteins were transferred to a 0.45 μm nitrocellulose filter (Thermo Scientific) using the Mini Blot Module (Invitrogen) at 10 V for 1 hour and 15 minutes at room temperature. After the transfer, non-specific binding to the filter was blocked overnight at 4°C in blocking buffer (binding buffer containing 3% BSA), and then the filter was probed in blocking buffer for 1 hour with streptavidin-HRP (1:10,000 dilution) at room temperature. After the incubation, the membrane was washed four times (10 min each) with binding buffer, followed by detection of bound Vip3Aa with enhanced chemiluminescence substrate (Super Signal West Pico PLUS, Thermo Scientific) and imaged in an Amersham Imager 600 (GE Healthcare). The intensity of the bound Vip3Aa band signal was measured by densitometry using the ImageJ software.^62^

Saturation binding assays with Vip3Aa labeled with Alexa-fluor were performed by incubating BBMV proteins (15 μg) with increasing amounts of labeled Vip3Aa (0.07 μg, 0.15 μg, 0.3 μg, 0.6 μg, 1.2 μg,1.8 μg, and 2.4 μg) with or without a 100-fold molar excess of unlabeled Vip3Aa in a final volume of 100 µL of binding buffer. The samples were incubated at 25°C for two hours and then centrifuged at 15,000 rpm for 15 minutes at room temperature. The supernatant was removed, and the pellet was resuspended in 500 µL of ice-cold binding buffer and centrifuged again. The final pellet was solubilized in 20 µL of measuring buffer (0.125 M Tris, 0.14 M SDS, 20% glycerol) and then applied to individual wells of a 1/2 Area OptiPlate-96 (Revvity), and the fluorescence at 488 nm measured in a Biotek Synergy H1 microplate reader. A standard curve using known amounts of fluorescence-labeled Vip3Aa was constructed to estimate the amounts of bound toxin represented by the fluorescence detected in binding reactions.

Binding saturation assays with radiolabeled Vip3Aa were performed as described previously,^37^ except that Tween-20 was not included in the binding buffer as tests with Tween-20 yielded the same qualitative observations but reduced bound toxin signal (data not shown). Reactions to determine total binding included 20 μg of BBMV and increasing amounts of Vip3Aa-I^125^(5 nM, 10 nM, 30 nM, 50 nM, and 100 nM) in a final volume of 0.1 mL of binding buffer (137 mM NaCl, 10 mM Na_2_HPO_4_, 2.7 mM KCl, 1.8 mM KH_2_PO_4_, 0.1%BSA, pH 7.4). Non-specific binding was determined by including a 300-fold molar excess of unlabeled Vip3Aa. After 1 hour of incubation at room temperature, reactions were stopped by centrifugation and washed as for assays with fluorescently labeled toxin. Two independent assays using different BBMV and radiolabeled toxin preparations were performed in two technical replicates for each input toxin concentration.

## 3 RESULTS

### 3.1 Transcriptome differences associated with resistance to Vip3Aa

Individual RNASeq library sizes and alignment statistics to the reference genome are presented in the Supporting Information (Tables S2-S3). A principal component analysis (PCA) plot of the samples sequenced showed variability among LA-RR samples, while LA-SS samples clustered together (Supporting Information, Fig. S3). A total of 2,900 differentially expressed genes (DEGs) were detected when comparing transcriptomes between LA-SS and LA-RR strains (Log2 fold change > 1, adjusted P < 0.05), with 1,300 downregulated and 1,600 upregulated genes in LA-RR relative to LA-SS. This DEG list was condensed after curating for DEGs that showed consistent expression trends among all six replicates sequenced per strain to 294 downregulated and 246 upregulated genes. The annotation of functional groups by GhostKoala in KEGG did not provide clarity to identify a pathway associated with the mechanism of resistance in the LA-RR strain (Supporting Information Table S4). A list of the top 10 upregulated and downregulated genes in LA-RR compared to LA-SS is presented in Supporting Information Table S5. Table 1 presents a list of selected DEGs encoding enzymes potentially involved in Vip3Aa mode of action, detoxification enzymes, and predicted membrane proteins (as annotated by RefSeq).

**Table 1.** Constitutively differentially expressed genes (DEGs) between 4^th^ instar susceptible (LA-SS) and Vip3Aa-resistant (LA-RR) FAW larvae. Only DEGs with consistent differences in expression in all six biological replicates, predicted to be involved in detoxification, Vip3Aa mode of action, or predicted membrane proteins with domains exposed to the extracellular cell surface, are included.

| Gene annotation | Gene accession | Log2Fold change | SEM | P-Adj value |
| --- | --- | --- | --- | --- |
| Chymotrypsin-1-like | XM_050694314.1 | -10.21 | 1.17 | 1.31E-15 |
| Multidrug resistance protein homolog 49, ABCB | XM_035579610.2 | -2.71 | 0.65 | 3.27E-05 |
| Trypsin alkaline-C like | XM_035596935.2 | -3.43 | 0.70 | 1.09E-06 |
| Chitin synthase-CHS-2 like | XM_050696839.1/XM_050696840.1 | -1.96 | 0.66 | 0.014 |
| Membrane alanyl aminopeptidase | XM_035583073.2 | -3.22 | 0.76 | 2.28E-05 |
| TP53-regulated inhibitor of apoptosis | XM_035583648.2 | -2.01 | 0.37 | 5.76E-08 |
| Facilitated trehalose receptor tret-1 like | XM_050704759.1 | -2.92 | 0.62 | 2.35 E-06 |
| Glutathione S transferase -2 like | XM_050693842.1 | -6.13 | 1.12 | 4.76E-05 |
| Plasma membrane calcium transporting ATPase-2 | XM_050699533.1 | -1.93 | 0.35 | 4.47E-08 |
| Membrane-bound alkaline phosphatase | XM_035584271.2 | 4.04 | 0.87 | 6.85E-05 |

None of the genes encoding proteins previously reported as putative Vip3Aa receptors or involved in resistance to FAW were identified as DEGs, except the chitin synthase-2 gene (GenBank accession: XM_050696839.1), which had 3.89-fold reduced expression in LA-RR compared to LA-SS (Table 1). Downregulation of chitin synthase-2 was confirmed by RT q-PCR (Supporting Information, Fig. S4). While no direct comparisons were performed before the collapse of the LA-SS strain, no significant differences in midgut chitin amounts were detected when comparing LA-RR to an alternative reference susceptible FAW strain (Benzon) (Fig. 1A).

**Figure 1.**
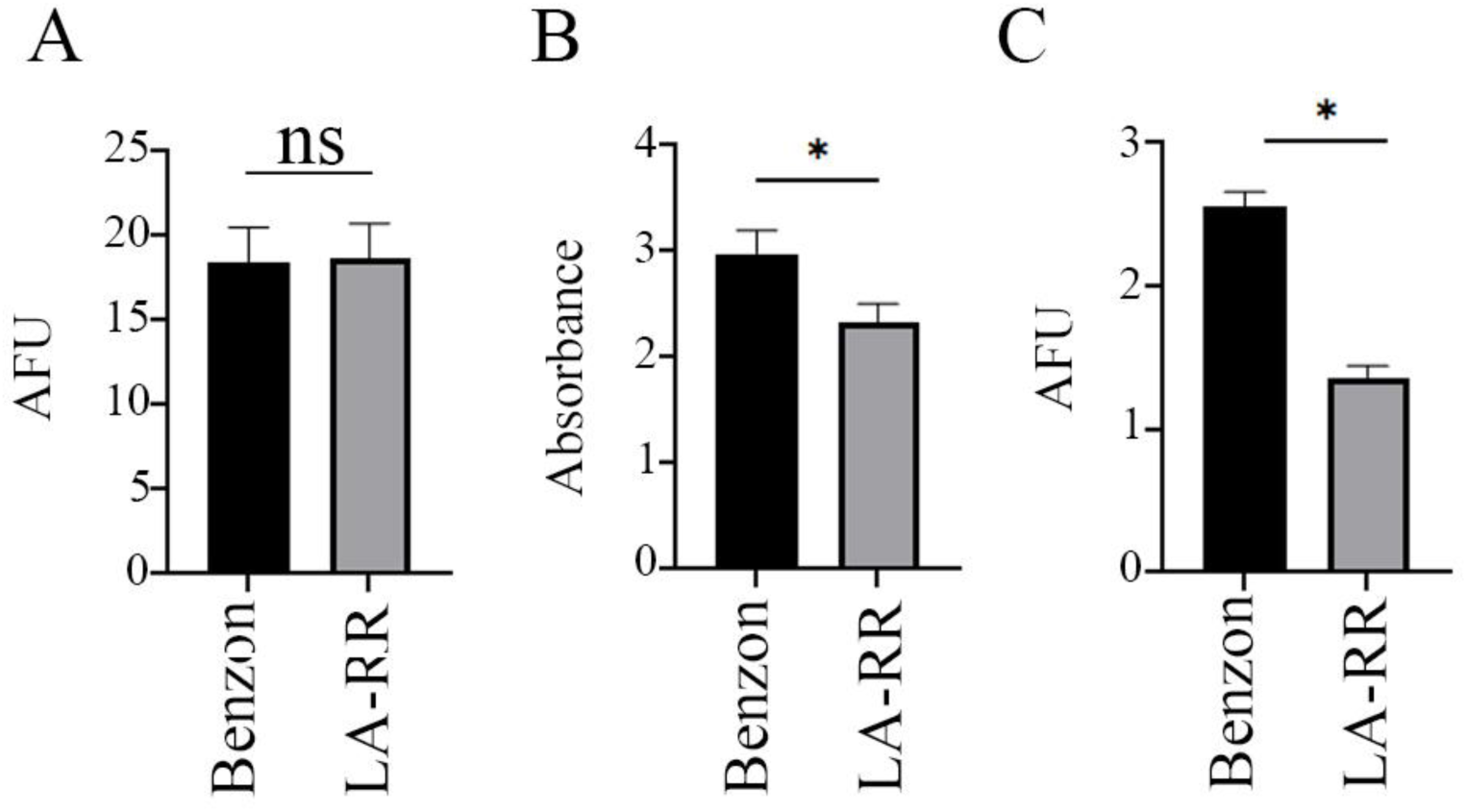
Estimation of midgut chitin content (A), and trypsin (B), and chymotrypsin (C) activities in midgut fluids from susceptible (Benzon) and Vip3Aa-resistant (LA-RR) strains of FAW. (A) The chitin content in 4^th^ instar larval midguts was quantified by calcofluor staining and measuring the fluorescence at 355 nm excitation. Shown are the means and corresponding standard error (SEM) calculated from 12 individual midguts for each strain. The significance of differences was tested by pooled t-test (P < 0.05). NS = not significant, AFU = Arbitrary fluorescent unit (raw fluorescent reads/10^4^). The relative difference in the activity of trypsin (B) and chymotrypsin (C) enzymes in 10 μg of gut fluid proteins were measured using L-BAPNA and Suc-Ala-Ala-Pro-Phe-AMC as substrates, respectively. Shown are the means and corresponding standard error (SEM) calculated from three biological replicates with two technical replicates each. Significant differences between strains (unpaired t test, P < 0.05) are indicated by an asterisk.

Specific chymotrypsin (GenBank accession XM_050694314.1) and trypsin (XM_035596935.2) genes were consistently down-regulated in LA-RR compared to LA-SS (Table 1), and this was further confirmed by RT-qPCR (Supporting Information, Fig. S4). While testing of specific serine protease activity in gut fluids from LA-SS was not possible due to the unexpected collapse of that colony, significantly reduced specific activities of trypsin (unpaired t-test P = 0.032) and chymotrypsin (unpaired t-test, P = 0.0001) were detected in gut fluids from LA-RR compared to larvae from the alternative reference susceptible colony (Benzon) (Figs. 1B and 1C).

A membrane-bound alkaline phosphatase (GenBank accession XM_035584271.2) homologous to a protein previously reported at reduced levels in midguts of Vip3Aa-resistant *Heliothis virescens*^63^ was also among the DEGs (Table 1). However, in LA-RR this alkaline phosphatase was upregulated 16.4-fold, translating into higher alkaline phosphatase activity in midgut brush border membrane vesicles from LA-RR compared to LA-SS (data not shown).

The list of curated DEGs also included members of protein families involved in the mode of action of Cry proteins (multidrug resistance protein homologous to ABCB transporters and membrane alanyl aminopeptidase), an inhibitor of apoptosis, and calcium and trehalose transporters with unknown participation in the Vip3Aa mode of action. A glutathione S transferase gene involved in detoxification was substantially down-regulated in LA-RR compared to susceptible larvae.

### 3.2 Effect of reduced serine protease activity on Vip3Aa protoxin processing

Drastically slower processing of Vip3Aa protoxin was observed when using gut fluids from LA-RR compared to LA-SS (Fig. 2A) and Benzon (Fig. 2B) larvae. A band of the expected size for Vip3Aa toxin was observed after 30 minutes of incubation with gut fluids from LA-SS or Benzon larvae, and its relative abundance continued to increase over time. In contrast, Vip3Aa protoxin processing by the LA-RR gut fluids was drastically slower, and no Vip3Aa toxin band was markedly visible even after 120 min of incubation. Differences in Vip3Aa processing over time were quantified using densitometry (Supporting Information, Fig S5A). These differences in Vip3Aa processing were not observed when comparing the processing of Cry1F protoxin (Supporting Information, Figs. S1 and S5B), which occurred for gut fluids from both LA-SS and LA-RR within 30 minutes.

**Figure 2.**
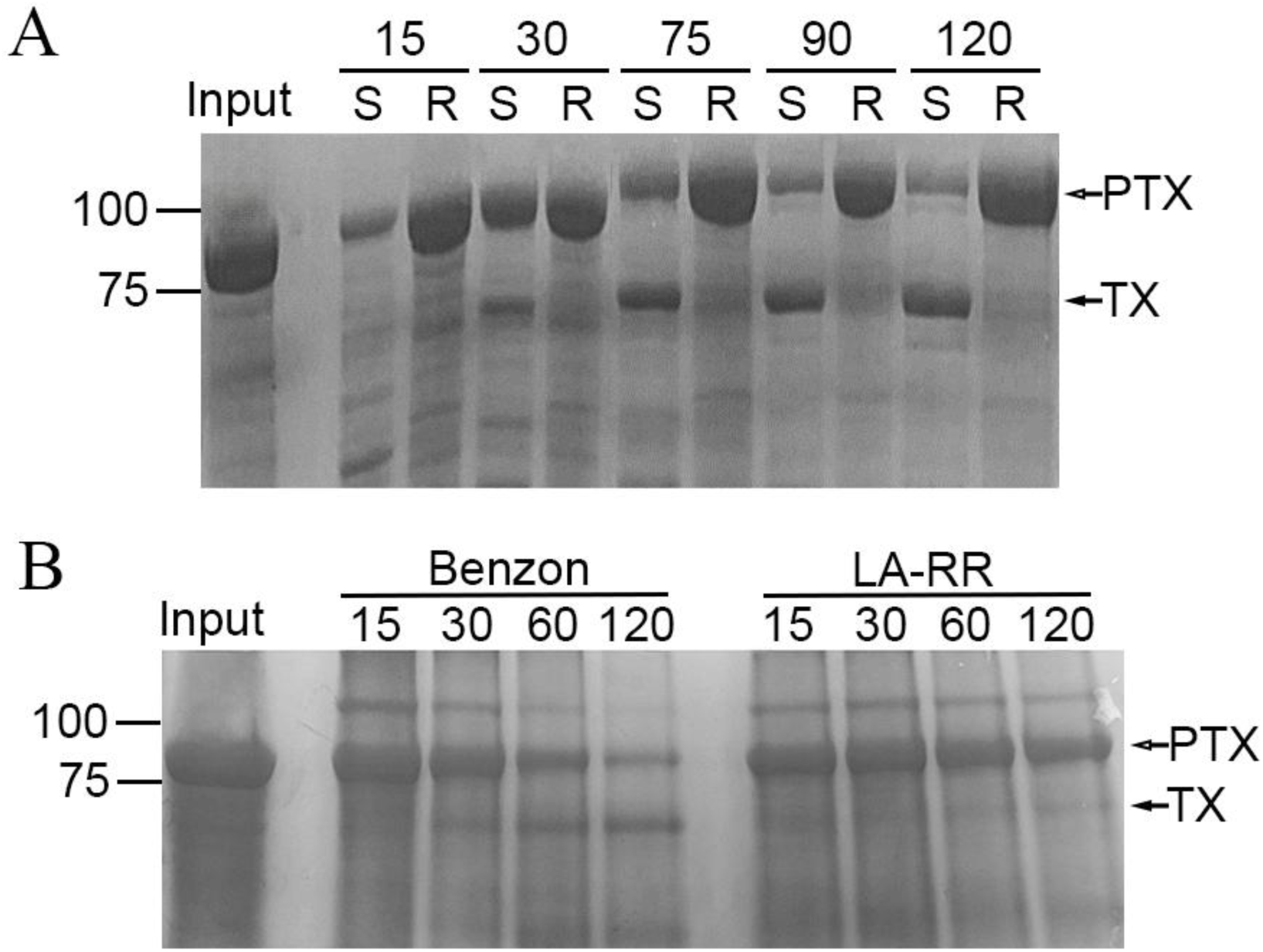
Processing of Vip3Aa protoxin by midgut fluids from larvae of susceptible (LA-SS and Benzon) and Vip3Aa-resistant (LA-RR) strains of FAW. Purified Vip3Aa protoxin (80 µg, I) was incubated with 0.4 µg of gut fluid proteins from the LA-SS (A), Benzon (B), and LA-RR (A and B) strains. After different lengths of incubation, as indicated in the figure, processing was visualized with SDS-10%PAGE and staining for total protein. The experiment was repeated thrice (A) or twice (B) with independent biological replicates with similar results. (PTX= protoxin, TX= toxin)

### 3.3 Slower Vip3Aa protoxin processing reduces midgut damage

Disruption of gut homeostasis by intoxication with Bt proteins results in a gut pH drop,^64,65^ which can be used as a proxy to monitor Cry1Ac and Vip3Aa damage to the midgut epithelium.^37^ Under control conditions, the average pH value in the foregut, midgut, and hindgut of Vip3Aa-susceptible (Benzon) and LA-RR larvae did not significantly differ (paired t-test, P = 0.231). Treatment with Vip3Aa protoxin or toxin did not affect the pH of foregut and hindgut in Benzon or LA-RR larvae (one-way ANOVA followed by Tukey post-hoc test, P = 0.519 for LA-RR, P = 0.161 for Benzon). On the other hand, treatment with Vip3Aa protoxin or toxin significantly reduced the midgut pH in Benzon larvae from 9.75 ± 0.15 (mean ± standard error) to 7.3 ± 0.13 and 7.4 ± 0.12, respectively (one-way ANOVA followed by Tukey post hoc test, P < 0.05) (Fig. 3A). In contrast, the midgut pH in LA-RR larvae was only significantly reduced (one-way ANOVA followed by Tukey post-hoc test, P < 0.05) from 9.13 ± 0.27 to 6.92 ± 0.12 when larvae were exposed to activated Vip3Aa toxin, with no significant changes when using protoxin (P = 0.966) (Fig. 3B).

**Figure 3.**
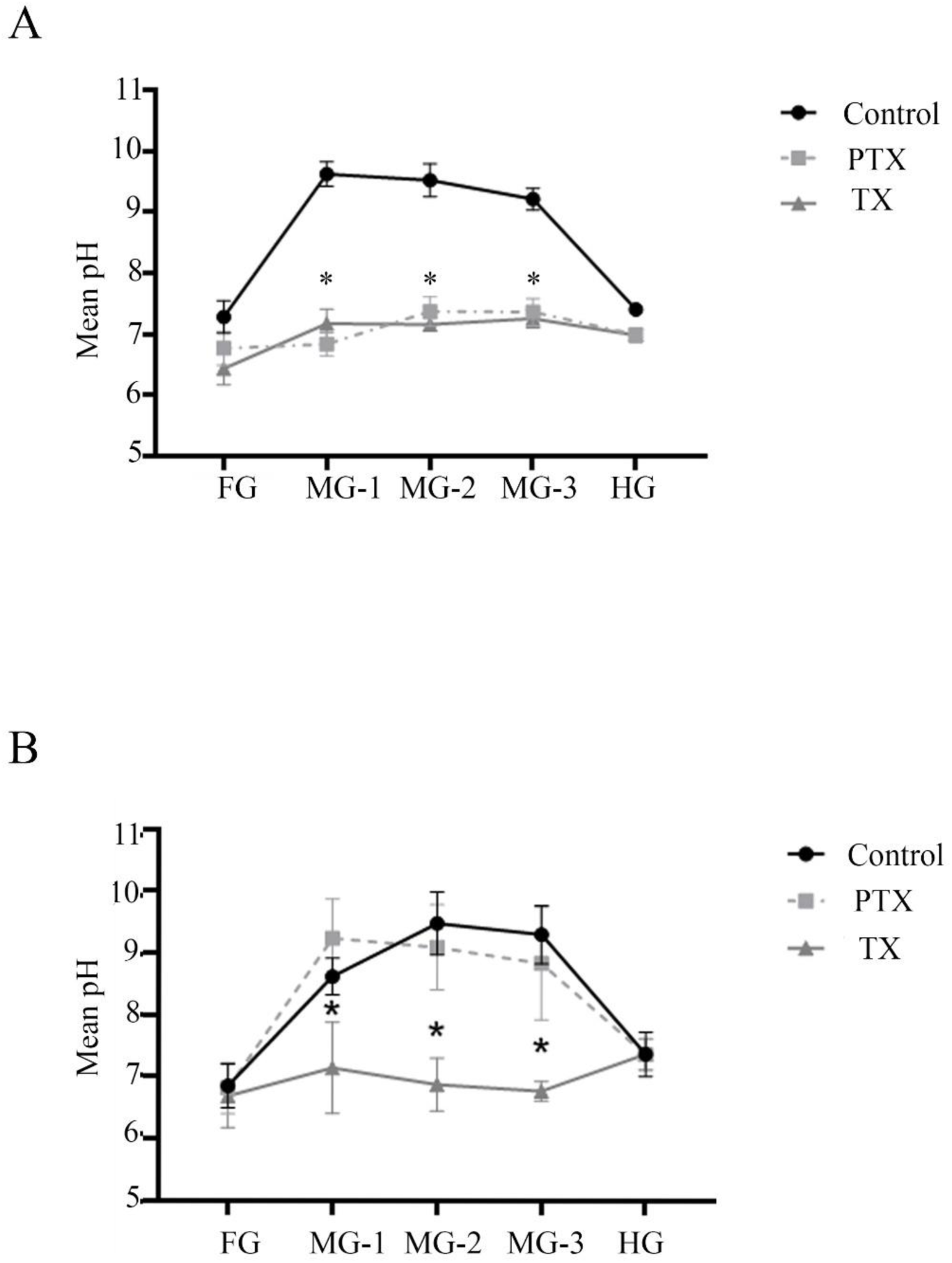
Change in gut pH in FAW larvae from the Benzon (A) and LA-RR (B) strains after feeding for 2 hours on a sucrose solution (control) or the same diet containing Vip3Aa protoxin (PTX) or activated toxin (TX). Larvae were dissected and the pH in five consecutive gut regions (from anterior to posterior: foregut, midgut 1, midgut 2, midgut 3, and hindgut) was measured using a microelectrode. Shown are the mean and corresponding standard error (SEM) from six larvae (biological replicates). An asterisk denotes statistically significant differences (one-way ANOVA, Tukey HSD, P < 0.05) among the different diets. FG= Foregut, MG= Midgut, HG= Hindgut.

### 3.4 Contribution of slower Vip3Aa protoxin processing on resistance

In testing the hypothesis that reduced rates of Vip3Aa processing contribute to resistance, we performed bioassays of LA-RR neonates exposed to a discriminatory dose of Vip3Aa protoxin^19^ pre-processed with gut fluids from LA-RR or Benzon larvae (Supplementary Information Fig. S6). In these bioassays, bypassing the slower processing of LA-RR by pre-processing with midgut fluids from Benzon resulted in significantly higher toxicity (unpaired t-test, P = 0.002) compared to when pre-processing with gut fluids from LA-RR larvae (Fig. 4A). Higher LA-RR susceptibility to Vip3Aa pre-processed by Benzon was also reflected in a significantly lower (unpaired t-test, P = 0.001) average mass of surviving larvae (Fig. 4B).

**Figure 4.**
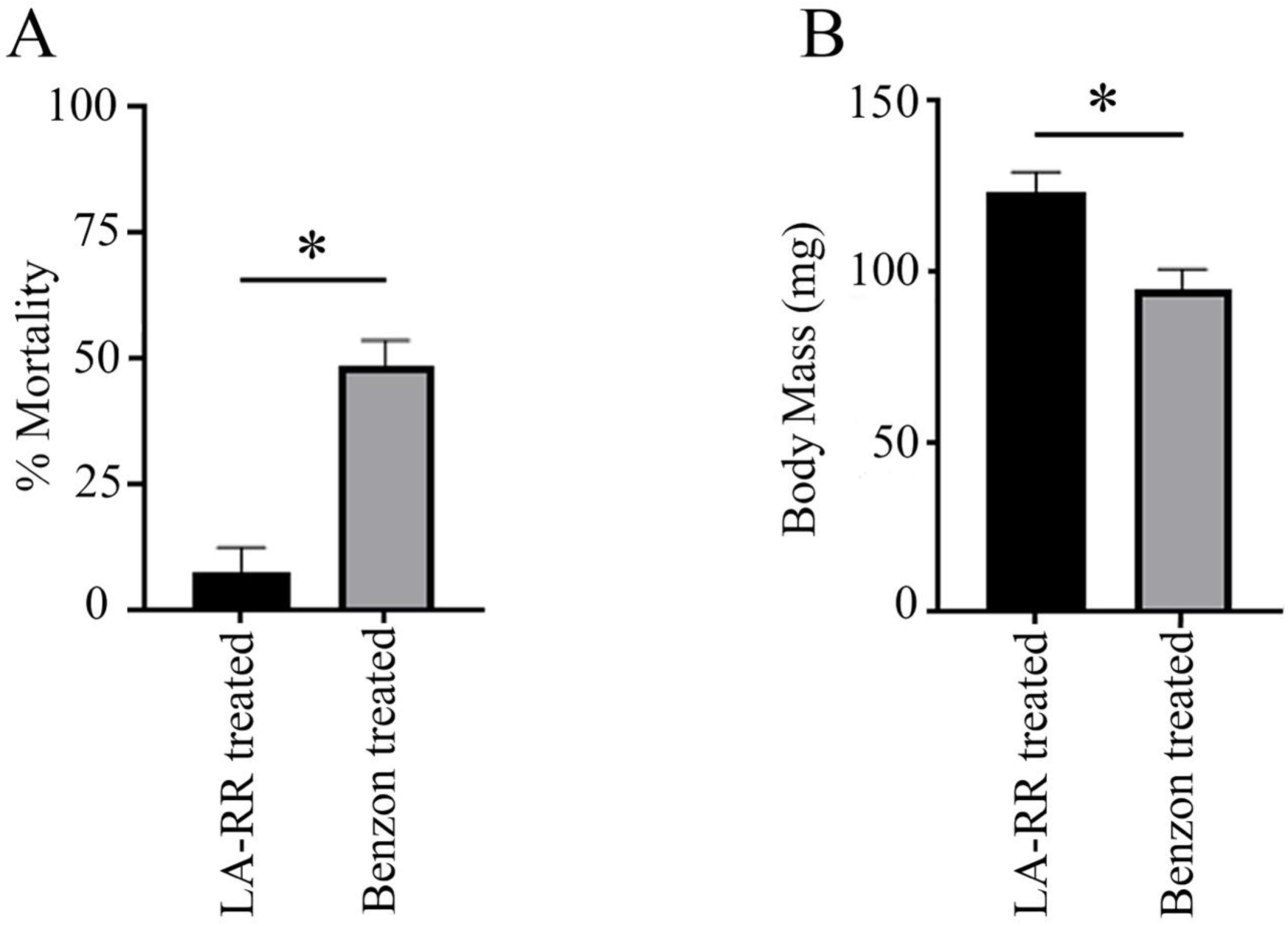
Mean percentage mortality (A) and average body weight of survivors (B) of LA-RR larvae nine days after feeding on a meridic diet treated with a discriminatory dose of Vip3Aa protoxin (5 μg/cm^2^) pre-processed with gut fluids from larvae of the Benzon or LA-RR strains of FAW. Shown are the mean and corresponding standard error (SEM) from a total of 64 larvae tested for each treatment. Significant differences (unpaired t-test, P < 0.05) are indicated by an asterisk.

### 3.5 Vip3Aa binding assays

Binding assays with Vip3Aa toxin labeled and detected with three different methods (Alexa-fluor 488, biotin, and I-125 radioisotope) detected specific binding to BBMV from all the tested strains (Fig. 5). Specific Vip3Aa binding did not saturate for the range of input labeled toxin concentrations tested, preventing accurate estimations of binding parameters. When using fluorescent-labeled Vip3Aa, there was a tendency for higher specific binding in BBMV from LA-SS and Benzon compared to LA-RR, which was only significant for the three highest toxin concentrations tested (one-way ANOVA, Tukey HSD, P = 0.002 for LA-SS and P = 0.013 for Benzon) (Fig. 5A).

**Figure 5.**
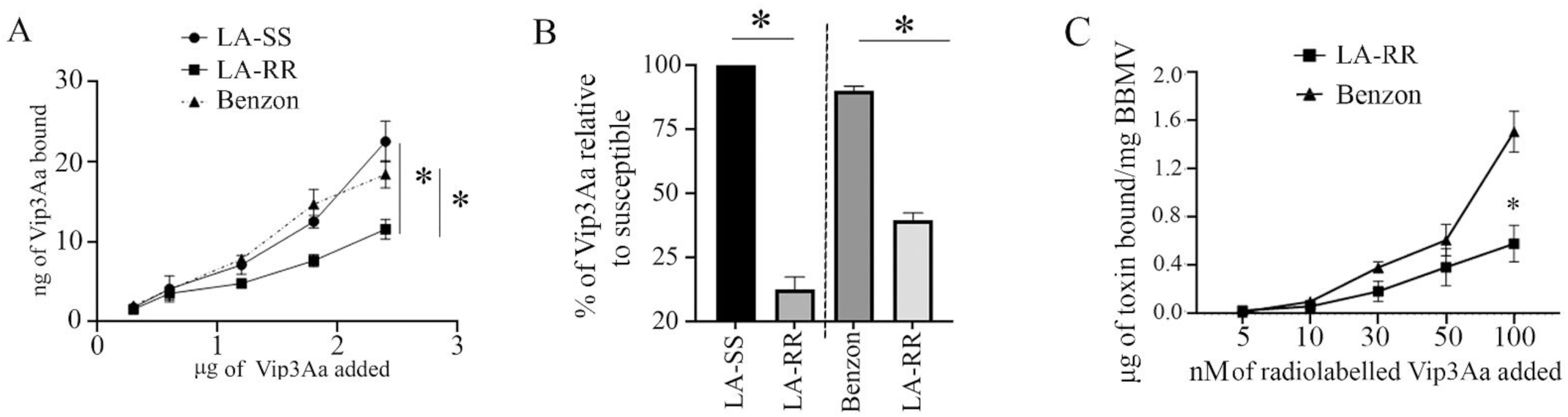
Binding of Vip3Aa toxin to midgut brush border membrane vesicles (BBMV) from susceptible (LA-SS and Benzon) and Vip3Aa-resistant (LA-RR) strains of FAW as tested with three different methods. (A) Specific binding of increasing amounts of Alexa-fluor labeled Vip3Aa to BBMV (15 μg) from the LA-SS, Benzon, and LA-RR strains. Reactions including 100-fold molar excess of unlabeled Vip3Aa were used to estimate non-specific binding. The bound fluorescent toxin was recovered by centrifugation, and its fluorescent signal was compared to a standard curve to estimate the amounts of bound labeled toxin. Non-specific binding was subtracted from total binding (without competitor) to calculate the specific binding shown in the graph. Each point and standard error bar are representative of three biological replicates. Significant specific binding differences (one-way ANOVA, Tukey HSD, P < 0.05) compared to the LA-RR strain are indicated with an asterisk. (B) Quantification by densitometry of biotin-labeled Vip3Aa (0.3 μg) specific binding to BBMV (25 μg) from the LA-SS, Benzon and LA-RR strains. After binding reactions, BBMV pellets and bound Vip3Aa were resolved by SDS-10%PAGE and transferred to a nitrocellulose filter. Densitometry was performed on the detected biotinylated toxin bands (see Fig. S7 for an example). Binding in the presence of a 100-fold excess of unlabeled Vip3Aa was used to estimate non-specific binding, which was subtracted from total binding (without competitor) to estimate specific binding. The bars represent the mean percentage and corresponding standard error from three biological replicates (different BBMV preparations). Specific binding in LA-RR was normalized considering the total binding for LA-SS and Benzon as 100%. Due to strain availability, experiments comparing binding between LA-SS and Benzon with LA-RR were performed in separate experiments, the respective LA-RR bars resulting from each comparison are shown. Asterisks denote a statistically significant difference (unpaired t-test, P < 0.05). (C) Specific binding of increasing amounts of Vip3Aa-I^125^ to BBMV (20 μg) from the Benzon and LA-RR strains. Reactions including 300-fold molar excess of unlabeled Vip3Aa were used to estimate non-specific binding. The bound radiolabeled toxin was recovered by centrifugation, and non-specific binding was subtracted from total binding (without competitor) to calculate the specific binding shown in the graph. Each point and standard error bar are representative of two biological replicates (different BBMV and Vip3Aa-I^125^) each performed in duplicate. Significant specific binding differences were only detected for the highest ligand concentration tested (unpaired t-test, P = 0.006), as indicated with an asterisk.

Western blots comparing biotinylated Vip3Aa binding to BBMV from LA-SS and Benzon with LA-RR were performed at different times depending on strain availability. Quantification by densitometry of the chemiluminescent signal from bound biotinylated Vip3Aa (Supporting Information, Fig. S7) in the absence (total binding) or presence of 100-fold excess unlabeled Vip3Aa (non-specific binding) allowed estimation of specific (total minus non-specific) binding. Specific biotinylated Vip3Aa binding was highest to BBMV from the LA-SS and Benzon strains, and was significantly reduced in BBMV from LA-RR larvae compared to BBMV from LA-SS (paired t-test, P = 0.003) and Benzon (paired t-test, P = 0.009) strains (Fig. 5B).

Comparison of the specific Vip3Aa-I^125^ binding was only possible between BBMV from the LA-RR and Benzon strains due to the collapse of LA-SS. These tests detected a clear tendency for higher specific binding to BBMV from Benzon compared to LA-RR in all the tested Vip3Aa-I^125^ concentrations (Fig. 5C), but was only statistically significant for the highest Vip3Aa-I^125^ concentration tested (unpaired t-test, P = 0.006).

## 4 DISCUSSION

Despite evidence supporting that resistance to Vip3Aa may be emerging in the Western Hemisphere,^14,19,20^ there is no mechanistic data available on resistance to Vip3Aa in FAW populations from this native range of FAW. We provide evidence supporting that reduced Vip3Aa protoxin processing, through down-regulation of specific serine protease genes, and reduced Vip3Aa toxin binding are both associated with the Vip3Aa resistance phenotype in the LA-RR FAW strain from Louisiana (USA). Transcriptome profiling of LA-RR did not detect the down-regulation of the *SfMyb* transcription factor previously reported as linked with resistance to Vip3Aa in an FAW strain from China.^66^ However, we cannot rule out the possibility that genes trans-regulated by the *SfMyb* transcription factor may still contribute to the resistant phenotype in LA-RR. Expression of chitin synthase-2 was reduced in LA-RR compared to susceptible larvae, as described for an FAW strain from China.^67^ However, reduced expression of chitin synthase-2 in LA-RR larvae did not translate into a significant reduction of midgut chitin content compared to Benzon. These observations suggest distinct resistance mechanisms to Vip3Aa may occur in strains from the native and invasive range of FAW. However, further molecular details are needed to determine if the distinct observations for Vip3A-resistant strains from the invasive and native range of FAW represent alterations at different stages of the same pathway. Moreover, RNA-seq data from our study did not detect altered expression of the *SfVipR1* gene (XM_035599784.2), recently proposed as relevant to Vip3Aa intoxication,^38^ in LA-RR.

Unfortunately, the unexpected collapse of the LA-SS strain prevented us from using this strain in further experiments. Based on the rationale that mechanistic differences in the Vip3Aa mode of action related to resistance would remain consistent between LA-RR and any susceptible strain, we used the Benzon strain as an alternative susceptible reference. Comparisons of Vip3Aa protoxin processing and toxin binding between LA-SS and Benzon did not detect relevant differences, supporting the use of Benzon as an appropriate reference susceptible strain for comparisons with LA-RR. Differences in genomic background between Benzon and LA-RR could have complicated genetic analyses, yet these were outside the scope of the current study.

While the current paradigm advocates for alterations in midgut receptors hindering toxin binding as the most common mechanism of resistance to insecticidal Bt proteins in lepidopteran pests, altered protoxin processing in resistant strains has also been reported.^68^ The few mechanistic studies available commonly report a reduced rate of protoxin processing in Vip3Aa-resistant lepidopterans.^35,36^ Importantly, reduced processing and binding are not necessarily mutually exclusive, and there are examples of both mechanisms contributing to a resistant phenotype to Cry toxins.^69^ However, this study represents the first example of this phenomenon in a strain resistant to Vip3Aa.

The monogenic inheritance of resistance reported for the LA-RR strain^19^ suggests that a single locus may be capable of simultaneously generating reduced processing and binding phenotypes. There is sufficient evidence in the literature supporting this hypothesis. For instance, retrotransposon-mediated disruption of a signaling pathway trans-regulates the expression of multiple midgut receptor and non-receptor genes and explains resistance to Cry1Ac in *Plutella xylostella.*^70,71^ In our transcriptome profiling, we detected multiple MAPK pathways altered in the LA-RR compared to the LA-SS strain (see DEGs full list), which could be involved in the resistance phenotype. Previous work on brown planthopper (*Nilaparvata lugens)* identified Krüppel-like (KNRL) transcription factors as capable of trans-regulating the expression of trypsin-like and chymotrypsin-like enzymes as well as other genes.^72^ GATA transcription factors regulate trypsin expression in the mosquito *Anopheles gambiae*^73^ and *Drosophila*.^74^ Importantly, GATAe was identified as the only GATA factor regulating the expression of Cry receptors (cadherin, ABCC2, and alkaline phosphatase) in midgut cells of *Helicoverpa armigera*.^75^ Moreover, genes possibly trans-regulated by *Sfmyb*, the transcription factor altered in a Vip3Aa-resistant strain of FAW from China, include a serine protease and other proteins that could represent receptors for Vip3Aa.^22^ Further work is needed to identify the locus responsible for the Vip3Aa resistance phenotype in LA-RR.

Reduced Cry protoxin processing in resistant lepidopteran insects is commonly due to reduced midgut protease activity.^76^ While pH is critical for Vip3Aa toxin binding to the midgut epithelium^77^, midgut pH did not differ between LA-RR and Vip3Aa-susceptible larvae under control conditions. Therefore, the detected reduced expression and activity of trypsin and chymotrypsin genes could explain the slower rate of Vip3Aa protoxin processing in LA-RR. Similarly, significantly reduced serine protease enzyme activity associated with slower Cry protoxin processing was reported in a Bt-resistant strain of *Plodia interpunctella*^57,78^ and Cry1Ac-resistant strains of *H. virescens*.^69^ The increased susceptibility of LA-RR larvae when using Vip3Aa protoxin pre-processed by midgut fluids from susceptible larvae highlights the relevance of altered processing for resistance. This observation is in line with the critical role of Vip3Aa protoxin processing in determining toxicity in *Spodoptera* spp.^79,80^ However, pre-processing did not completely restore susceptibility in LA-RR, in line with the proposed participation of reduced Vip3Aa toxin binding in the resistant phenotype. This lack of susceptibility may appear in conflict with the similar drop in midgut pH observed after resistant and susceptible larvae fed on trypsin-activated Vip3Aa. However, it is important to consider the different amounts of toxin and larval instar used in these tests, which hinders direct correlation of results. Alternatively, it is possible that the amount of midgut damage sustained, while inducing similar midgut pH drop, may differ between the two strains.

In general, Vip3Aa processing involves proteolytic cleavage of a 20-kDa region from the C terminus by serine proteases.^31,32,79^ In FAW, Vip3Aa protoxin processing is primarily performed by cationic trypsin and anionic chymotrypsin.^79^ Only one chymotrypsin-1-like and trypsin alkaline type -C genes were consistently down-regulated in LA-RR compared to susceptible larvae, supporting their importance in Vip3Aa processing. Remarkably, the processing of Cry1F was not affected in LA-RR relative to LA-SS, suggesting that the down-regulated serine proteases are specific to Vip3Aa but do not affect Cry protein processing. This hypothesis is supported by the lack of cross-resistance to Cry1F in LA-RR.^19,42^ Reduced activity of trypsin-like and chymotrypsin-like enzymes in LA-RR is in agreement with the transcriptomic data and provides a strong association between reduced trypsin and chymotrypsin activities, processing, and Vip3Aa resistance. Since fully processed Vip3Aa is needed to effectively bind to receptors in midgut cells,^80^ reduced Vip3Aa processing probably reduces the amount of protein able to effectively bind to midgut receptors. The detected reduced binding of fully processed Vip3Aa to BBMV from LA-RR compared to susceptible FAW strains would provide a second line of defense against intoxication. This binding reduction was observed using three different toxin labeling and detection methods, each with specific advantages and limitations. More gentle conditions in biotinylation and fluorescence labeling probably better preserve Vip3Aa functionality. Western blots with biotinylated Vip3Aa also detected the bound labeled protein after electrophoretic separation, allowing for the detection of potential processing. On the other hand, experiments with radiolabeled protein had the highest sensitivity and measured bound Vip3Aa directly on the BBMV.

Multiple putative Vip3Aa receptors have been proposed from *in vitro* studies with cell cultures, including prohibitin-2, fibroblast growth factor receptor, and scavenger receptor-c.^81-83^ However, none of these putative Vip3Aa receptors were confirmed in functional assays with FAW larvae^84^ or had altered expression in the LA-RR strain. A gene encoding a protein homologous to a membrane-bound alkaline phosphatase in Vip3Aa-resistant *H. virescens* had increased expression in LA-RR. However, this protein did not function as a Vip3Aa receptor when expressed in cultured insect cells.^63^ Further research is needed to determine the mechanism and receptors involved in the reduced Vip3Aa binding in LA-RR.

## 5 CONCLUSION

The experimental evidence presented in this study supports that resistance to Vip3Aa in the LA-RR strain is likely mediated by both slower Vip3Aa processing from reduced expression of trypsin and chymotrypsin-like protease genes and reduced specific binding of the activated toxin to unidentified receptors in the midgut epithelium. This study presents the first mechanistic description of resistance to Vip3Aa in a strain from the native FAW range and provides evidence for the first example of multiple mechanisms associated with resistance to Vip3Aa.

## Supporting information

see DEGs full list

## ACKNOWLEDGEMENTS

This work was supported by the Agriculture and Food Research Initiative program, project award no. 2021-67013-33567, from the U.S. Department of Agriculture’s National Institute of Food and Agriculture.

## CONFLICT OF INTEREST DECLARATION

The authors declare that they have no known competing financial interests or personal relationships that could have appeared to influence the work reported in this paper.

## DATA, MATERIALS and SOFTWARE AVAILABILITY

The raw transcript sequences reported in this paper have been deposited in the National Center for Biotechnology Information Sequence Read Archive (NCBI SRA) database as accession no. PRJNA1196488. Additional raw data generated in this study are included in the Raw Data Appendix file.

## Notes

### Competing Interest Statement

The authors have declared no competing interest.

