## Supporting Information, Figs. S1 for "Reduced processing and toxin binding associated with resistance to Vip3Aa in a resistant strain of fall armyworm (*Spodoptera frugiperda*) from Louisiana"

**Figure S1-** Processing of Cry1F protoxin by midgut fluids from larvae of the LA-SS and LA-RR strains of fall armyworm (FAW, *Spodoptera frugiperda*). Partially purified protoxin (80 µg) was incubated with 0.4 µg of gut fluid proteins from each strain. The reactions were stopped at different time points and samples were resolved by SDS-10%PAGE and visualized using a stain for total protein (ProtoBlue Safe). The experiment was repeated twice with independent biological replicates with similar results. S= LA-SS, R= LA-RR, PTX- Input partially purified protoxin, TX- Cry1F toxin band)

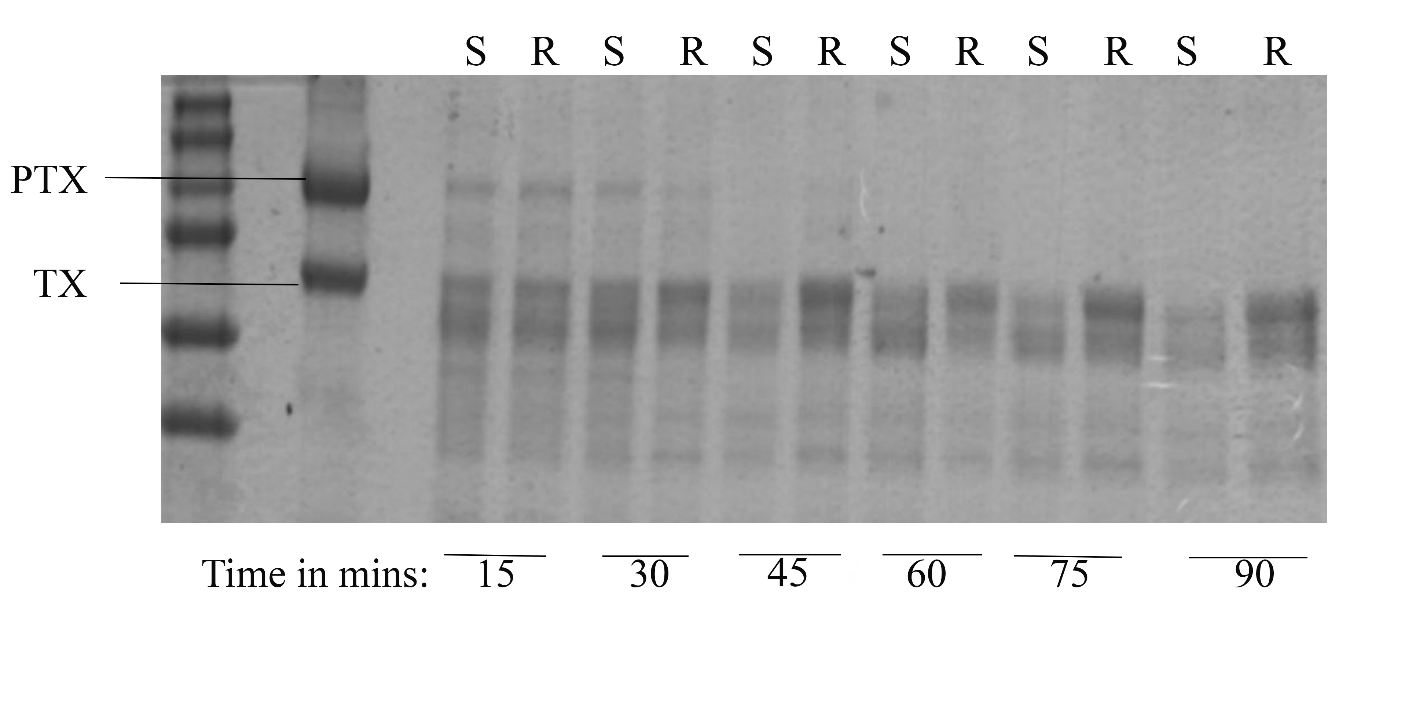

**Figure S2** – Testing successful labeling of Vip3Aa toxin with Alexa fluor 488. A) Purified Vip3Aa toxin labeled with Alexa fluor 488 was resolved by SDS-10%PAGE and then visualized in a ChemiDoc MP imaging system. I-1: 10 ng of input labeled Vip3Aa, I-2: 1 ng of input labeled Vip3Aa, I-3: 2 ng of input labeled Vip3Aa, I-4: 4 ng of input labeled Vip3Aa. B) Detection of Alexa-fluor 488 labeled Vip3Aa (0.6 μg) bound to midgut brush border membrane vesicles from larvae of the LA-SS strain of FAW. The binding assay was performed as described in Materials and Methods without (total binding) or with (non-specific binding) a 100-fold molar excess of unlabeled Vip3Aa toxin. 1- LA-SS total binding, 2- LA-SS non-specific binding.

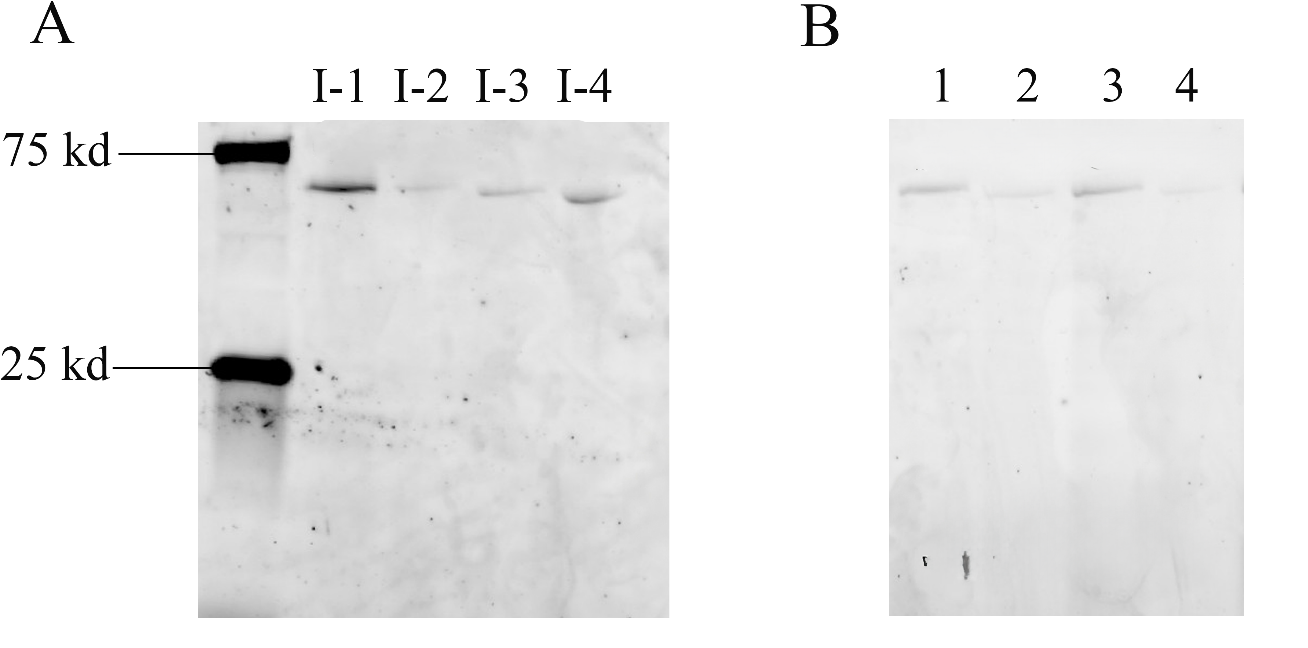

**Figure S3** – Principal component analysis (PCA) plot of differentially expressed genes (DEGs) in six individual midguts (each) from larvae of the LA-RR and LA-SS strains of FAW. Transcriptomic data was analyzed with DE-seq2 to generate the PCA plot.

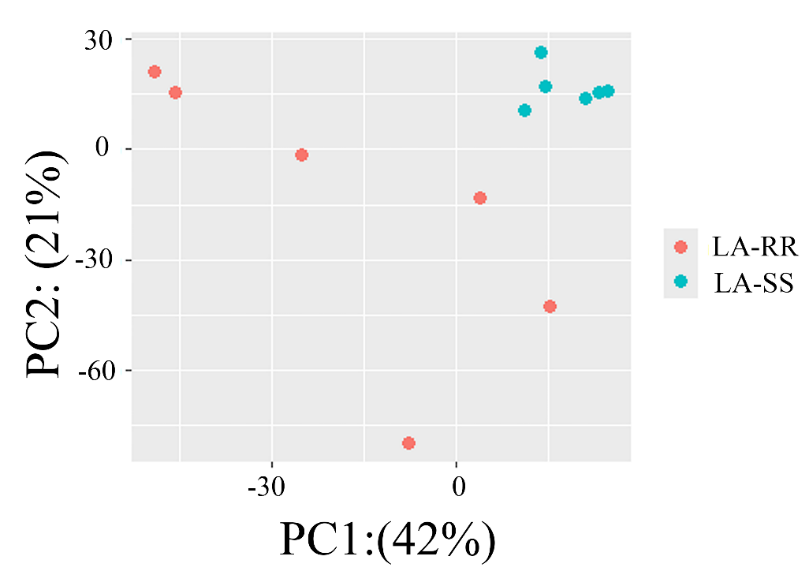

**Table S1:** Primer efficiencies for primers used for quantitative PCR shown in Fig. S4, determined using a 10-fold serial dilution of cDNA (1:10 to 1:1000), with each dilution run in technical duplicates due to template limitations. All qPCR reactions produced a single melting curve peak, confirming specificity and absence of non-specific amplification. The primer sequences are listed 5’ to 3’ direction

| Gene Name | Forward primer | Reverse primer | Primer efficiency |
| --- | --- | --- | --- |
| Trypsin Alkaline-C | GGGGTCATACTGCATACTACGG | AGTGACCCCTAAGACCCCAG | 105.35% |
| Chitin Synthase-2 | ACCTTCCGGGGGATCATACA | ACACAAACGTGAAACCTACTCA | 100.98% |
| Chymotrypsin-1 like | ATTGGCAGCTTTGCCTTTCC | GAGGACAGTGCGTTGAGTGA | 97.43% |

**Table S2**: Raw read data from the Illumina library preparation of an RNAseq library with RNA from six individual midguts each of a susceptible (LA-SS) and a Vip3Aa-resistant (LA-RR) strain of FAW. Effective (%) = Total percentage number of possible reads minus error (Error %) and no signal reads. Q20 (%) = Percentage of bases with a quality score of 20 or higher, denoting an error rate of 1 in 100. Q30 (%) = Percentage of bases with a quality score of 30 or higher, denoting an error rate of 1 in 1000. GC (%) = Percentage GC content.

| Strain | Sample | Raw reads | Effective (%) | Error (%) | Q20 (%) | Q30 (%) | GC (%) |
| --- | --- | --- | --- | --- | --- | --- | --- |
| LA-RR | D1 | 172210518 | 94.08 | 0.02 | 98.75 | 95.79 | 40.62 |
|  | D2 | 210831198 | 97.56 | 0.02 | 98.54 | 95.69 | 48.63 |
|  | D3 | 189444222 | 96.85 | 0.02 | 98.35 | 95.10 | 44.36 |
|  | D4 | 177106246 | 97.55 | 0.02 | 98.61 | 95.69 | 48.56 |
|  | D5 | 214574790 | 98.04 | 0.02 | 98.64 | 95.81 | 48.08 |
|  | D6 | 157762702 | 96.56 | 0.02 | 98.72 | 95.79 | 42.90 |
| LA-SS | E1 | 207152160 | 97.97 | 0.02 | 98.65 | 95.65 | 47.08 |
|  | E2 | 196821988 | 98.01 | 0.02 | 98.64 | 95.69 | 47.59 |
|  | E4 | 156064940 | 98.64 | 0.02 | 98.58 | 95.46 | 47.04 |
|  | E5 | 189044302 | 97.91 | 0.02 | 98.73 | 95.87 | 46.41 |
|  | E8 | 162813930 | 96.68 | 0.02 | 98.79 | 96.17 | 47.04 |
|  | E9 | 177861724 | 97.92 | 0.02 | 98.71 | 96.00 | 48.20 |

**Table S3**: Alignment statistics of raw reads from the samples in Table S2 to the reference FAW genome (accession no: GCF_023101765.2) . Shown are percentages of unique matches (Unique), matches to multiple loci (Multimap), not mapped to the genome (Mismatch), too short for mapping (Unmapped short), or unmapped for any other reason (Unmapped other).

| Sample | Unique | Multimap | Mismatch | Unmapped short | Unmapped other |
| --- | --- | --- | --- | --- | --- |
| D1 | 78.87 | 6.66 | 0 | 12.5 | 0.66 |
| D2 | 76.03 | 14.96 | 0 | 7.22 | 0.83 |
| D3 | 80.63 | 6.57 | 0 | 10.6 | 1.52 |
| D4 | 81.13 | 11.41 | 0 | 5.99 | 0.52 |
| D5 | 78.53 | 13.59 | 0 | 6.26 | 0.87 |
| D6 | 80.19 | 7.55 | 0 | 9.61 | 1.27 |
| E1 | 80.53 | 11.95 | 0 | 6.07 | 0.44 |
| E2 | 81.52 | 11.17 | 0 | 5.72 | 0.55 |
| E4 | 82.92 | 9.82 | 0 | 5.81 | 0.5 |
| E5 | 83.4 | 8.85 | 0 | 6.3 | 0.37 |
| E8 | 81.72 | 9.91 | 0 | 6.52 | 0.53 |
| E9 | 82.18 | 10.14 | 0 | 6.15 | 0.63 |

**Table S4:** Functional analysis of the upregulated and downregulated genes in LA-RR compared to LA-SS performed using GhostKoala in KEGG. The annotated functional groups and the number of genes in each group are presented. Genes that are present in multiple categories are counted multiple times

| Expression | Functional Group Annotation | Number of genes |
| --- | --- | --- |
| Up | Protein families: genetic information processing | 147 |
|  | Protein families: cell signaling and cellular processes | 120 |
|  | Environmental information processing | 102 |
|  | Cellular processes | 66 |
|  | Organismal systems | 58 |
|  | Genetic information processing | 52 |
|  | Protein families: metabolism | 46 |
|  | Carbohydrate metabolism | 44 |
|  | Lipid metabolism | 38 |
|  | Human diseases | 20 |
|  | Amino acid synthesis | 20 |
|  | Glycan biosynthesis and metabolism | 19 |
|  | Nucleotide biosynthesis | 15 |
|  | Unclassified protein families | 13 |
| Down | Protein families: genetic information processing | 172 |
|  | Genetic information processing | 128 |
|  | Carbohydrate metabolism | 83 |
|  | Energy metabolism | 81 |
|  | Protein families: signaling and cellular processes | 48 |
|  | Lipid metabolism | 48 |
|  | Environmental information processing | 38 |
|  | Cellular processing | 38 |
|  | Glycan biosynthesis | 30 |
|  | Organismal systems | 29 |
|  | Protein families:metabolism | 29 |
|  | Metabolism | 25 |
|  | Amino acid metabolism | 24 |
|  | Unclassified protein families | 24 |
|  | Nucleotide biosynthesis | 20 |

**Table S5:** List of the top ten genes upregulated and downregulated in larvae from the LA-RR compared to the LA-SS strains from the RNA-seq experiment. LfcSE = Log2 fold change standard error, EC = Expression change (down or up-regulated)

| Gene name | Normalized transcript count | Log2 Fold change | lfcSE | pAdj | Annotated function | EC |
| --- | --- | --- | --- | --- | --- | --- |
| LOC118275360 | 4744.576 | -11.94 | 1.21 | 1.01E-19 | uncharacterized LOC118275360 | Down |
| LOC118267671 | 393.3841 | -10.21 | 1.17 | 1.31E-15 | chymotrypsin-1-like |  |
| LOC118281432 | 161.8522 | -8.24 | 0.89 | 1.74E-17 | cuticle protein 19 |  |
| LOC118267884 | 1611.165 | -7.42 | 0.91 | 2.04E-13 | collagenase-like |  |
| LOC118263044 | 4802.965 | -7.38 | 1.25 | 2.70E-07 | para-nitrobenzyl esterase-like |  |
| LOC118267896 | 49.82662 | -7.18 | 1.03 | 7.58E-10 | odorant receptor 4-like |  |
| LOC126911117 | 307.5467 | -6.68 | 0.72 | 1.07E-17 | trichohyalin-like |  |
| LOC118261750 | 7.783289 | -6.40 | 1.52 | 0.000343 | odorant receptor 49b-like |  |
| LOC118276858 | 451.5907 | -6.39 | 0.67 | 2.64E-18 | uncharacterized LOC118276858 |  |
| LOC118274101 | 19735.87 | -6.29 | 0.80 | 1.52E-12 | uncharacterized LOC118274101 |  |
| LOC118272140 | 87141.33 | 9.79 | 0.87 | 1.14E-25 | uncharacterized LOC118272140 | Up |
| LOC118263615 | 240.3232 | 8.86 | 1.45 | 8.58E-08 | serine protease inhibitor dipetalogastin |  |
| LOC118275914 | 723.0623 | 8.73 | 0.74 | 5.00E-28 | L-threonine ammonia-lyase |  |
| LOC118268700 | 94.86209 | 8.72 | 1.16 | 1.60E-11 | putative nuclease HARBI1 |  |
| LOC118276098 | 102.5258 | 8.31 | 1.35 | 8.04E-08 | uncharacterized LOC118276098 |  |
| LOC118272399 | 41.37443 | 7.76 | 1.08 | 1.83E-10 | uncharacterized LOC118272399 |  |
| LOC118279246 | 43.59918 | 7.44 | 1.34 | 1.52E-06 | golgin subfamily A member 6-like protein 22 |  |
| LOC118269266 | 20.05089 | 6.71 | 1.32 | 1.26E-05 | degenerin-like protein asic-1 |  |
| LOC118278686 | 199.1495 | 6.68 | 1.31 | 1.15E-05 | uncharacterized LOC118278686 |  |
| LOC118271788 | 2655.983 | 6.47 | 0.82 | 1.52E-12 | mucin-2-like |  |

**Figure S4**- Relative expression difference for selected DEGs in the midgut of 5^th^ instar LA-SS and LA-RR larvae quantified by RT-qPCR. Columns bars are the mean and the corresponding standard errors, respectively, from four biological replicates tested in technical triplicates. Expression detected for LA-SS was considered as the level of “1” when normalizing expression of LA-RR samples. Asterisks denote significant expression differences between LA-SS and LA-RR for each gene (pooled t-test, P < 0.05). Chtr= Chymotrypsin-1 like (XM_050694314.1), Tryp= Trypsin alkaline-C like (XM_035596935.2) and Chs-2= Chitin Synthase-2 like (XM_050696839.1/XM_050696840.1)

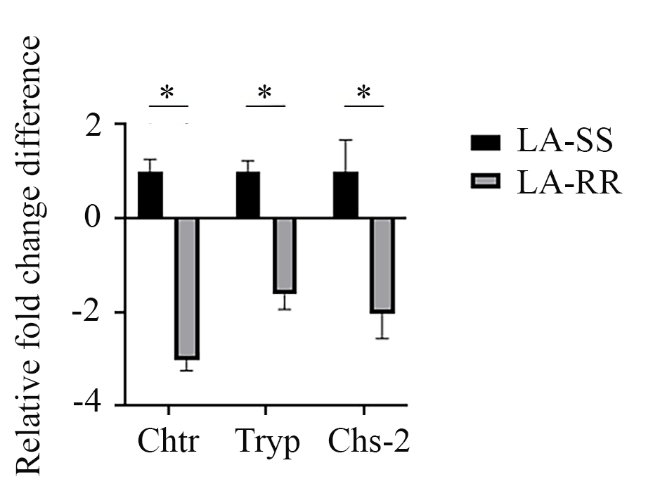

**Figure S5 –** Processing of Vip3Aa (A) and Cry1F (B) protoxin bands by midgut fluids from FAW larvae as quantified by densitometry. Each point and bar represent the mean and corresponding standard error from three biological replicates (different midgut fluid samples) for Vip3Aa and two for Cry1F.

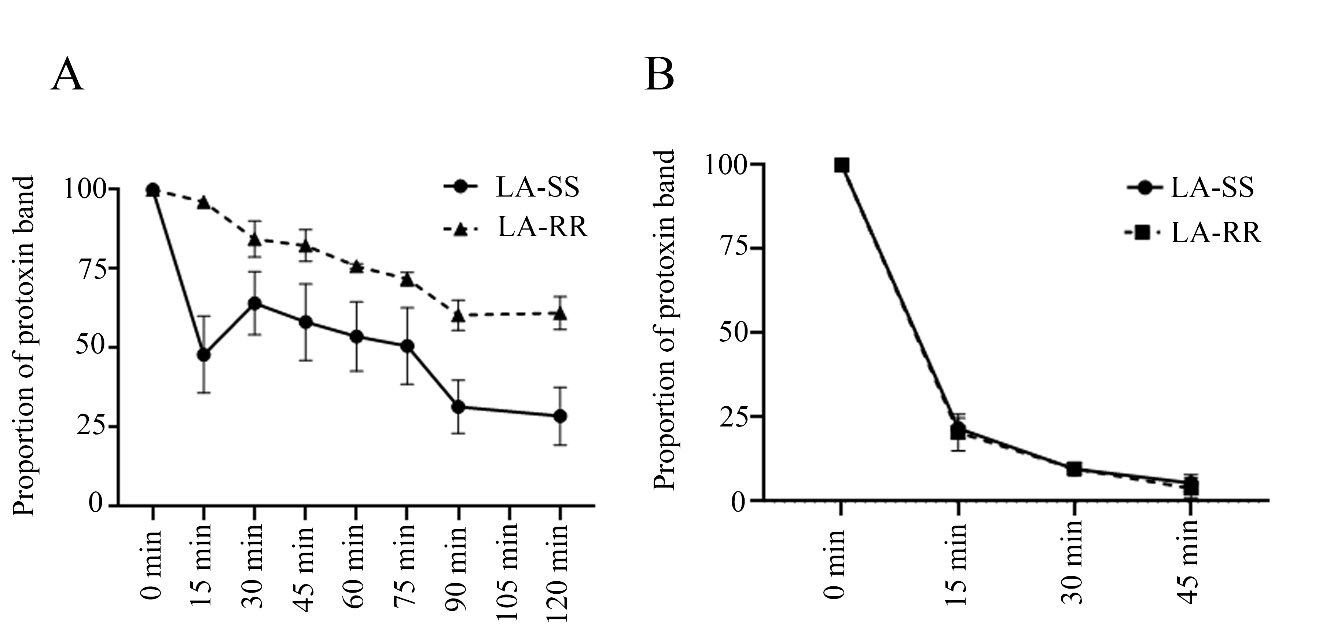

**Figure S6**- Diagram detailing the bioassay with the pre-processed Vip3Aa protoxin. The Vip3Aa39 protoxin was processed by gut fluids from LA-RR or Benzon larvae and then used on diet-overlay bioassays with LA-RR neonates to estimate toxicity. Shown are the 3D structures for Vip3Aa protoxin (also when processed by LA-RR gut fluids) and activated Vip3Aa toxin (when processed by Benzon gut fluids).

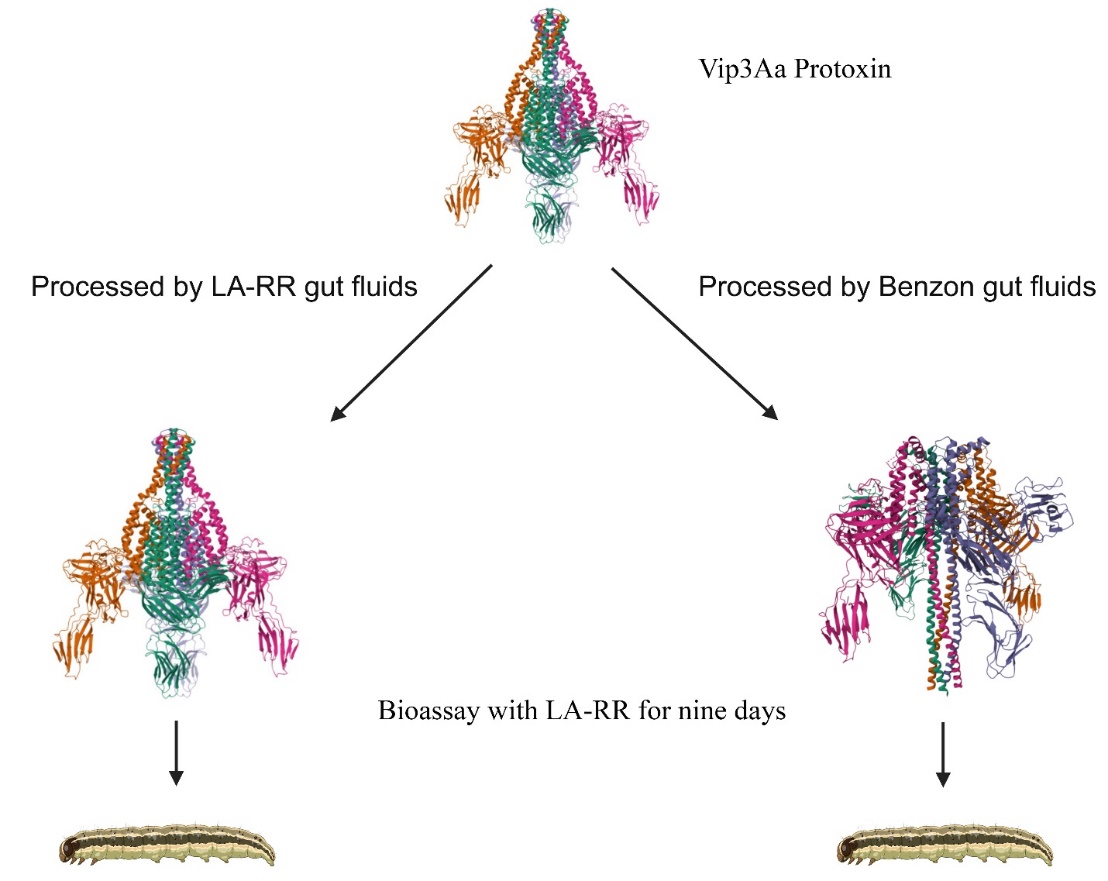

**Figure S7-** Detection of biotinylated Vip3Aa (0.3 μg) binding to midgut brush border membrane

vesicles (BBMVs) (25 μg) from the LA-RR, LA-SS, and Benzon, as indicated. BBMVs were incubated with biotinylated Vip3Aa without (total binding, T) or with a 100-fold molar excess of unlabeled Vip3Aa toxin (non-specific binding, NS). The bound toxin was recovered by centrifugation and washed twice with binding buffer before using streptavidin and enhanced chemiluminescence to detect the bands. Images within A) and B) are from the same filter and exposure length, lanes were cropped to eliminate lanes unrelated to the experiment. Vip3Aa = 10 ng of biotinylated Vip3Aa, used as a reference.

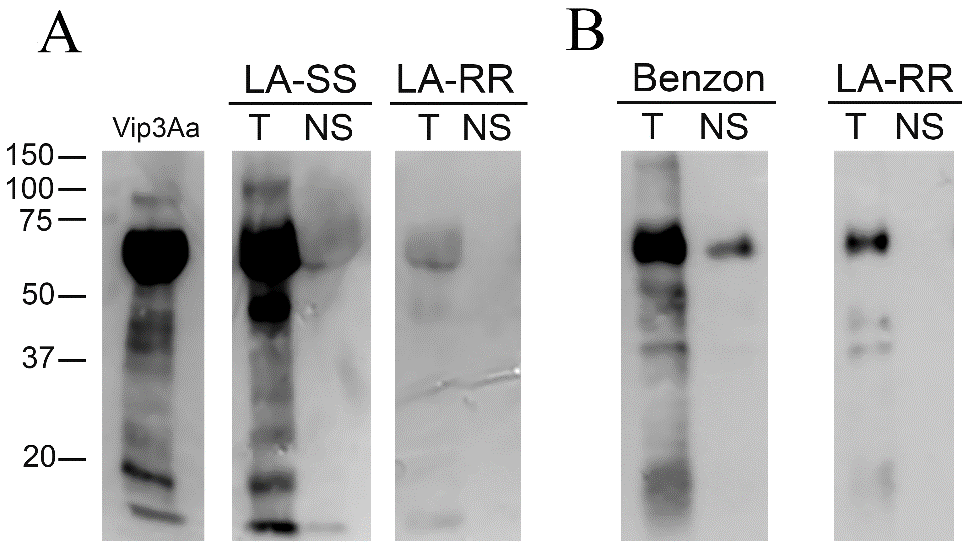
